# Chemogenetic control of GABAergic activity within the interpeduncular nucleus reveals dissociable behavioral components of the nicotine withdrawal phenotype

**DOI:** 10.1101/2024.07.05.602259

**Authors:** Anabel M. M. Miguelez Fernández, Shana Netherton, Seshadri B. Niladhuri, Patricia Rivera, Kuei Y. Tseng, Christian J. Peters

## Abstract

Chronic exposure to nicotine results in the development of a dependent state such that a withdrawal syndrome is elicited upon cessation of nicotine. The habenulo-interpeduncular (Hb-IPN) circuit contains a high concentration of nAChRs and has been identified as a main circuit involved in nicotine withdrawal. Here we investigated the contribution of two distinct subpopulations of IPN GABAergic neurons to nicotine withdrawal behaviors. Using a chemogenetic approach to specifically target Amigo1-expressing or Epyc-expressing neurons within the IPN, we found that activity of the Amigo1 and not the Epyc subpopulation of GABAergic neurons is critical for anxiety-like behaviors both in naïve mice and in those undergoing nicotine withdrawal. Moreover, data revealed that stimulation of Amigo1 neurons in nicotine-naïve mice elicits opposite effects on affective and somatic signs of withdrawal. Taken together, these results suggest that somatic and affective behaviors constitute dissociable components of the nicotine withdrawal phenotype and are likely supported by distinct subpopulations of neurons within the IPN.

## Introduction

Nicotine is a high-affinity agonist of nicotinic acetylcholine receptors (nAChRs), and chronic exposure results in the development of a dependent state such that a nicotine withdrawal syndrome is elicited upon cessation. Rodent models of nicotine withdrawal are characterized by behavioral changes that recapitulate symptoms of nicotine withdrawal described in humans (McLaughlin et al., 2015). The most common behavioral manifestations of nicotine withdrawal examined in rodents are increases in stereotypic behaviors referred to as somatic signs, and affective manifestations including signs of anhedonia, depression, hyperalgesia and, most commonly, anxiety (McLaughlin et al., 2015; Chellian et al., 2021).

Blocking nAChRs with an antagonist is sufficient to precipitate withdrawal in animal models of chronic nicotine exposure, which is strong evidence that nicotine withdrawal is mediated through plastic changes to cholinergic signaling (Malin et al., 1994; Damaj et al., 2003). Antagonizing cholinergic transmission by systemic injection of mecamylamine can precipitate withdrawal in mice chronically exposed to nicotine, and this phenotype can be phenocopied by injection of nAChR antagonists directly to the interpeduncular nucleus (IPN) (Salas et al., 2009). Block of IPN nAChRs is sufficient to precipitate somatic (Salas et al., 2009) or affective (Zhao-Shea et al., 2015) signs of nicotine withdrawal only in nicotine-exposed animals, underscoring the importance of this nucleus to the nicotine withdrawal phenotype. Knockout of α5 nAChR subunits in IPN suppression of somatic signs of nicotine withdrawal (Jackson et al., 2008). Optogenetic silencing of GAD2+ IPN neurons reduced somatic and affective signs in animals undergoing nicotine withdrawal (Klenowski et al., 2022). Taken together, these results suggest that plastic changes in IPN following chronic nicotine exposure, such as enhanced glutamate release and nAChR upregulation (Arvin et al., 2019), might drive the physical and affective manifestations of withdrawal.

A recent study has revealed that the IPN is composed of at least two major subpopulations of GABAergic neurons, both of which express the α5 nAChR subunit. These populations can be identified by their differential expression of the cell-surface adhesion proteins Epyc and Amigo1 (Ables et al., 2017). Epyc-IPN neurons are primarily local interneurons, while Amigo1-IPN neurons project to the Raphe nuclei and the laterodorsal tegmentum. Moreover, it has been shown that optogenetic silencing of Amigo1-IPN neurons prevents conditioned place preference for nicotine, whereas silencing Epyc-cells has no effect (Ables et al., 2017). Although both subpopulations receive major inputs from the medial habenula, this evidence suggests that they serve different functions in the circuit responsible for behavioral changes to nicotine. In this study, we used a chemogenetic approach to determine the role of these IPN subpopulations in driving behaviors associated with nicotine withdrawal in mice. Our results reveal separable contributions of the Amigo1 population to the somatic and affective phenotypes of withdrawal.

## Materials and Methods

### Animals

All experimental procedures were approved by the University of Illinois Chicago Institutional Animal Care Committee and met the National Institutes of Health guidelines for care and use of laboratory animals. Animals were housed in the University of Illinois Chicago Biologic Resources Laboratory in groups of 2-5 mice/cage under constant temperature (21-23°C) and light/dark cycle (14 hr/10 hr) with access to food and water *ad libitum.* Transgenic mouse lines were obtained from the Mutant Mouse Research Resource Center and originally generated by the GENSAT consortium (Ables et al., 2017) and were backcrossed to BL/6J using mice purchased from Jackson Laboratory. Mice were genotyped by PCR on genomic tail DNA using primers from the Washington University Genotyping Core Facility. Cre mouse lines were backcrossed at least 10 generations to BL/6J (Jackson stock no.000664) before these studies. Male and female mice 8 weeks or older were used in all experiments.

### Stereotaxic delivery of AAV-encoded DREADDs

Adeno-associated viral constructs AAV9-hsyn-DIO-mCherry, AAV9-hsyn-DIO-HM4D(Gi)-mCherry and AAV9-hsyn-DIO-HM3D(Gq)-mCherry were obtained from Addgene (viral prep #50459, #44362, # 44361 respectively; ≥ 1×10¹³ vg/mL titer) (Krashes et al., 2011) and kept at -80°C prior to use. On the day of the surgery, the virus was thawed and diluted 1:3 in sterile saline. Mice were anesthetized using >2% isoflurane (Somnosuite Unit, Kent Scientific), and dorsal skin was prepared by depilation and subdermal injection of 0.125% bupivacaine prior to mounting into a stereotaxic frame (Stoelting). Meloxicam (2 mg/kg, s.c) and ampicillin (10 mg/kg, s.c.) were used as analgesic and antibiotic, respectively. Vaporized isoflurane anesthesia (1.5-3.5%) was maintained throughout the procedure and body temperature kept at ∼37°C using a heating pad. After making a ∼1 cm incision along the midline of the scalp, and drilling a burr hole above the injection site, a 35G needle mounted to a 10 μl Nanofil syringe and driven by a UMPI perfusion pump (WPI) was advanced to the coordinates (in mm, relative to bregma): -3.400 A/P, 0.666 M/L and 4.854 D/V, at a lateral angle of 8°. 500 nL of viral solution was administered at a rate of 250 nL/min followed by a 5-minute rest period. The needle was withdrawn at a rate of 1 mm/minute. The incision was closed using wound clips. Animals were allowed to recover alone before returning to their home cage. A minimum of 2 weeks was allowed for AAV expression prior to experiments or additional surgery.

### Nicotine exposure

Mice were chronically exposed to nicotine by means of osmotic pump implanted via an incision between the scapulae that delivered a constant subcutaneous flow of 0.5μl/h for 14 days (RWD Bioscience). Pumps were filled with 0.9% saline for controls, or saline supplemented with nicotine at a concentration adjusted to individual body weight such that each mouse received nicotine at 1.0 g/kg/hour. (−)-Nicotine hydrogen tartrate salt was from Sigma-Aldrich.

### Behavioral procedures

Animals were acclimatized to the experimenter, the behavior room and to i.p. injections (using 0.9% saline) for 2 days before behavior testing. On behavior day, all mice were injected i.p. with 1 mg/kg Clozapine-N-Oxide and 3 mg/kg of mecamylamine (Damaj et al., 2003) at 30 min and 5 min before the start of the first test, respectively. Both drugs were obtained from Tocris. Next, they were placed in an elevated plus maze (EPM; Stoelting) to assess anxiety-related behavior. The apparatus consisted of a central platform (5 × 5 cm), two opposed open arms (35 × 5 cm) and two opposed closed arms with 15-cm-high dark walls, at a height of 50 cm. Mice were placed on the central platform facing an open arm and allowed to explore for 5 min while being recorded by an overhead camera. Time spent and distance traveled in each arm were obtained using EthoVision XT tracking software (Noldus). To account for time spent in the center zone (neither open nor closed), open arm time and distance indices were calculated as time or distance spent in open arms over the sum of open and closed arms.

Immediately after completion of the EPM test, mice were placed in a square arena (30 cm × 30 cm × 30 cm acrylic box with two opaque and two transparent sides) to examine physical signs of nicotine withdrawal for 15 min. Behavior was video-recorded and scored offline and blinded, using a custom-made push-button device for simultaneous recording of onset, offset and duration of events. Button presses were digitized as square waves in Axoscope and analyzed in Clampfit10 (Axon Instrument) using single-channel analysis mode. Signs were divided into the following categories: 1) grooming-related events including face or body scratching, genital licking or any part of the stereotyped sequential movements of the grooming pattern and 2) events indicating physical discomfort including head shaking, body twitching or jumping. In addition, the overhead video was analyzed using EthoVision XT to determine the distance travelled. Illumination for each behavior apparatus was approximately 200 lux, while the closed arms of the EPM led to lower brightness (∼80 lux).

### Histology

Following behavioral assays, mice were perfused with PBS followed by 4% paraformaldehyde under deep isoflurane anesthesia and brains were removed for verification of virus expression. Brains were post-fixed in 4% paraformaldehyde for 3-4hrs, washed in PBS and stored in 30% sucrose for cryoprotection. Brains were frozen, cryosectioned and directly mounted onto charged slides. Coronal 20μm sections were coverslipped using VectaShield mounting media with DAPI (Vector Laboratories) and visualized with a fluorescence microscope (Keyence BZ-X800).

### Ex-vivo electrophysiology

Independent groups of virus-injected mice were used to characterize the impact of DREADD activation on neuronal firing activity. Animals were perfused with ice-cold NMDG-based cutting solution (Ting et al., 2014) bubbled with 5% carbogen, containing (in mM): 93 NMDG, 2.5 KCl, 1.2 NaH2PO4, 20 HEPES, 25 glucose, 5 Na-ascorbate, 3 Na-pyruvate, 10 MgSO4, 0.5 CaCl2, pH adjusted to 7.3–7.4 with NMDG and osmolarity to 300–310 mOsm with mannitol. Brains were immediately removed to ice-cold, oxygenated NMDG cutting solution. Brains were sliced into 350 μm coronal sections using a VT 1000S vibratome (Leica Microsystems). Slices were placed in a holding chamber in NMDG-based cutting solution at 35°C for 15 min, then transferred to a chamber containing oxygenated HEPES-supplemented aCSF solution containing (in mM): 92 NaCl, 2.5 KCl, 1.2 NaH2PO4, 30 NaHCO3, 20 HEPES, 25 glucose, 5 Na-ascorbate, 3 Na-pyruvate, 2 MgSO4, 2 CaCl2, pH to 7.3–7.4 with HCl, osmolarity to 300–310 mOsm with mannitol, initially held at 35°C and allowed to cool to room temperature. Slices were held at room temperature for at least 45 min before recording.

Slices were continuously perfused with oxygenated aCSF using a peristaltic pump throughout recordings. Recording aCSF contained (in mM): 127 NaCl, 1.8 KCl, 26 NaHCO3, 12 KH2PO4, 1.3 MgSO4, 2.4 CaCl2, 15 glucose, with pH adjusted to 7.3–7.4 using HCl, and osmolarity to 304– 306 mOsm with mannitol. Pipette internal solution contained (in mM): 129.5 K-Gluconate, 6.5 KCl, 2 MgCl_2_, 4 Na-ATP, 0.4 Na-GTP, 0.2 EGTA, 10 Hepes, with pH adjusted to 7.2 using KOH and osmolarity adjusted to 295 with mannitol. Recording pipettes were pulled from filamented 1.5 OD, 0.86 ID borosilicate glass (Sutter Instrument) using a Sutter P97 puller and polished to a tip resistance of 4-8 MΩ using a Narishige microforge. Following establishment of a GΩ seal, whole cell mode was established using gentle suction, and cells were held for 2-5 minutes following patch rupture prior to recording. Rapid solution exchange used a Picospritzer 3 (Parker-Hanifin) to drive outflow from a 200 μm applicator held immediately above the neuron.

All recordings were done in Clampex10 using a MultiClamp 700B patch clamp amplifier and a Digidata 1550 digitizer (Axon Instruments) built on an Axioskop2FS stand (Carl Zeiss). Neurons were visualized with a CCD camera (QImaging) driven by μManager software. Electrophysiology data were analyzed using Clampfit10. Only cells with access resistance <35 mΩ were analyzed. To quantify passive properties, resting membrane potentials were measured in current clamp mode, after >20 s at 0 pA DC current injection. To quantify membrane excitability, DC current was injected to current clamped neurons to reach a rheobase value, which we defined as firing at a frequency of <0.1 Hz across >20 s; the resulting voltage at rheobase was compared before and after CNO application (2μM).

### Statistical Analysis

Data were summarized as mean ± SEM and differences among experimental conditions were considered statistically significant at *p*<0.05. One-way ANOVA was used as indicated in figure legends. Statistical analyses were performed with GraphPad Prism10.

## Results

To probe separable effects of activity in subpopulations of IPN GABAergic neurons on withdrawal behavior, we employed a chemogenetic approach targeted to two separate cre lines by targeting IPN neurons with AAV infection. We first examined the contribution of Amigo1-IPN neurons to behaviors associated with nicotine withdrawal using a chemogenetic method (see timeline in **Figure 1A**). Amigo1-cre mice (Ables et al., 2017) were infected in the IPN with hM4D(Gi), an inhibitory Designer Receptor Exclusively Activated by Designer Drugs (DREADD) by stereotaxic injection of AAV9-hSyn-DIO-hM4D(Gi)-mCherry (Krashes et al., 2011), which is an adeno-associated virus (AAV) that encodes cre-dependent hM4D(Gi) fused with an mCherry fluorophore. Control animals were sham injected with “empty vector” AAV9-hSyn-DIO-mCherry. The ability of hM4D(Gi) to manipulate Amigo1 neurons activity was tested using *ex vivo* brain slice electrophysiology from IPN neurons from hM4D(Gi)-mCherry positive neurons. IPN neurons were recorded under current clamp with 0 pA of direct current injection to quantify resting potential (-56.5 ± 1.8 mV, **Figure 1C** *left*). We then adjusted DC current in 10 pA steps for >20 s to a level where action potentials fired at <0.1 Hz, which we treated as “rheobase”, and quantified the voltage (-61.3 ± 3.2 mV, **Figure 1C** *right*) as a measure of excitability. Then, neurons were treated with CNO and allowed to equilibrate for >30 s. We then compared the resting potential (-63.3 ± 1.9 mV) and voltage at rheobase (-61.0 ± 3.7 mV) with values before CNO exposure. We observed a significant hyperpolarization of the resting membrane potential in the presence of CNO (p<0.0001, **Figure 1C** *left*), but no apparent change to voltage at rheobase (p=0.868, **Figure 1C** *right*). Next, we examined changes to affective behavior in response to nicotine by exposing Amigo1-hM4D(Gi) animals to nicotine or saline (vehicle) using a subcutaneous osmotic pump for 14 days. Following the 14 days exposure period, affective signs of withdrawal were evaluated using an elevated plus maze (EPM). DREADDs were activated by intraperitoneal injection of CNO (3 mg/kg) 30 minutes prior to the test, and mCherry controls were injected with CNO to control for any off-target effects of CNO. Nicotine withdrawal was precipitated by an i.p. injection of the nAChRs antagonist mecamylamine 5 minutes prior to behavior test. Open arm avoidance and increased occupancy of the closed arms of the maze were interpreted as reflecting heightened anxiety in the context of nicotine. We found that inhibition of Amigo1-IPN neurons prior to precipitating nicotine withdrawal reduced anxiety-like behaviors in animals chronically exposed to nicotine as evidenced by an increased open arm exploration (time 68.23±13.98 s vs. 203.5±24.64 s, p=0.0001, **Figure 1D**; index 0.28±0.06 vs. 0.75±0.08, p=0.0001, **Figure 1F**) and reduced closed arm occupancy (183.60±20.19 s vs. 66.88±23.51 s p=0.0003, **Figure 1E**) with respect to their mCherry counterparts. This reduction of anxiety-like behavior to the levels of nicotine-naïve animals was not explained by changes in locomotion activity (8.73±0.57 m vs. 5.74±0.64 m total distance traveled, p=0.0053, **Figure 1G**), since the distance travelled in the open arms was significantly larger than that in the closed arm, resulting in an open arm distance index undistinguishable from saline controls (0.60±0.08 vs. 0.73±0.09, p=4694, **Figure 1H**). Virus expression was confirmed in the IPN of all mice *post-mortem* by fluorescence microscopy (e.g.: **Figure 1B**). Next, using the same strategy (**Figure 2A, B**) we examined the contribution of a second population of neurons, marked by the gene *Epyc,* to affective behaviors. Epyc-cre mice were infected and prepared similar to Amigo1-cre animals and tested in the elevated plus maze, using similar mCherry sham infection and acute CNO i.p. administration to all animals as controls. Animals that had been exposed to nicotine displayed a typical pattern of enhanced anxiety evidenced by the reduced time spent (time 159.70±21.41 s vs. 80.50±22.73 s, p=0.0235, **Figure 2C**; index 0.60±0.07 vs. 0.30±0.08, p=0.0108 **Figure 2E**) and distance travelled in the open arms of the maze (index 0.53±0.07 vs. 0.24±0.05, p=0.0030, **Figure 2G**) compared to saline exposed controls. However, inhibition of Epyc-IPN neurons failed to alter the behavior of nicotine-exposed mice (open arm time 80.50±22.73 s vs. 101.90±15.23 s, p=0.7726, **Figure 2C**; closed arm time 190.50±22.52 s vs. 139.80±18.43 s, p=01814, **Figure 2D**; open arm index 0.30±0.08 vs. 0.44±0.07, p=0.3947 **Figure 2E**) or the distance travelled in the maze (7.99±0.77 m vs. 7.10±0.63 m vs. 8.54±0.65 m, **Figure 2F**) after precipitated withdrawal. Taken together, these results show that Amigo1 (and not Epyc)-IPN neuron activity is required for anxiety-like behaviors in nicotine-exposed mice after precipitated withdrawal.

**Figure 1.**
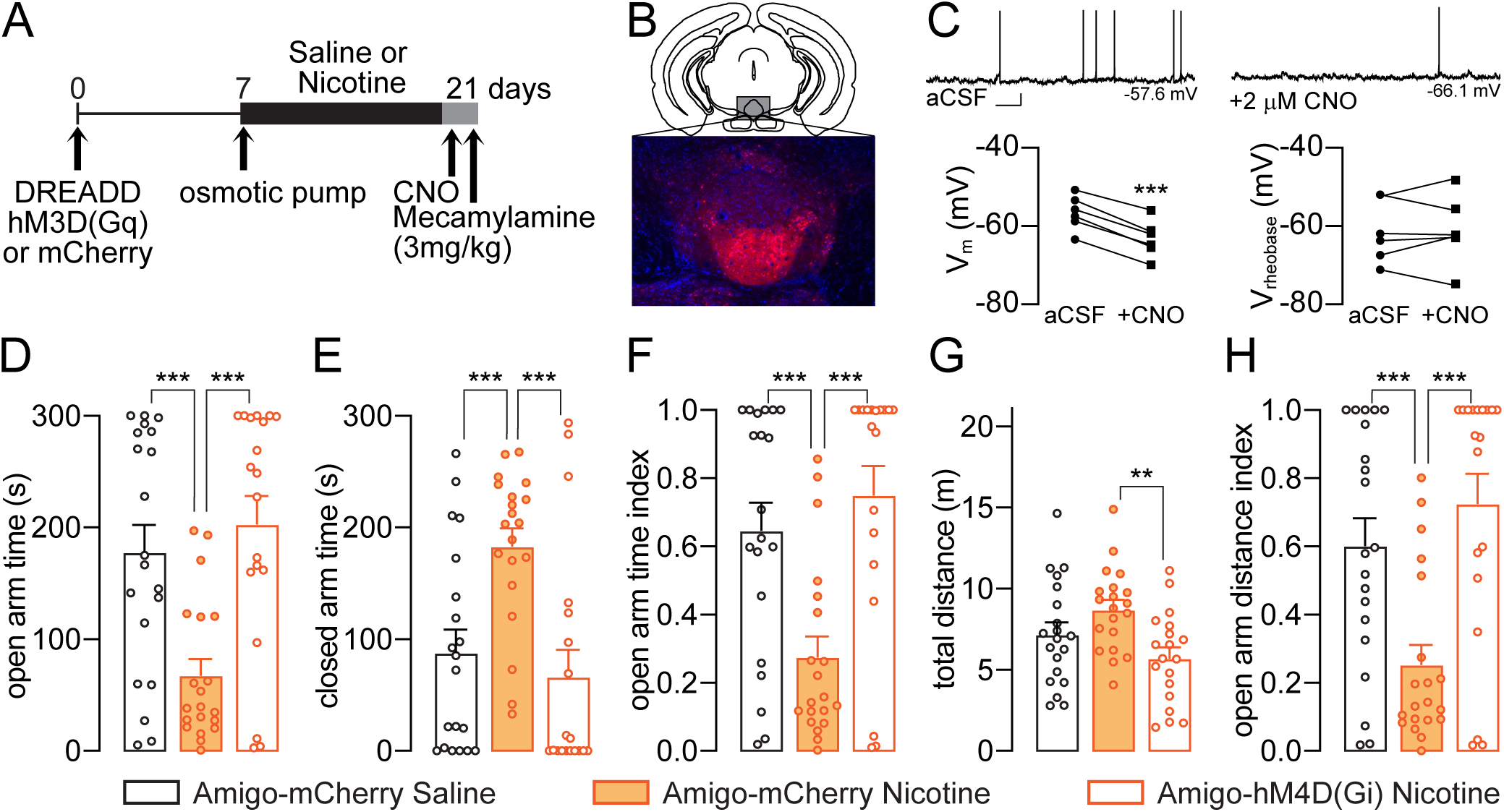
Inhibition of Amigo1-IPN neurons reduces anxiety-like behaviors in mice undergoing nicotine withdrawal. **A)** Timeline of the experimental design. Experimental groups are as follows: mCherry Saline (N=20), mCherry Nicotine (N=20), hM4D(Gi) Nicotine (N=19). **B)** Schematic diagram of virus injection target and representative image of virus expression in the IPN of Amigo1-cre male and female adult mice. **C)** *Top*: Sample traces of action potentials from Amigo1-cre cells expressing hM4D(Gi) in response to aCSF alone or supplemented with 2μM CNO. Scale bar indicates 10mV and 500ms and applies to all traces. *Left*: CNO application leads to a negative shift in resting membrane voltage compared with aCSF alone (n=6, t-test). *Right*: Rheobase voltage was determined by adjusting DC current to a level where action potentials fired at <0.1 Hz as another measure of excitability (n=6, t-test). **D-H)** Elevated plus maze. Maze exploration was assessed by evaluating time spent in the open (**D:** F_(2, 56)=_11.46, p<0.0001) and closed (**E:** F_(2, 56)_=9.804, p=0.0002) arms as well as the total distance travelled in the maze (**G:** F_(2, 56)_=5.327, p=0.0076) as a measure of locomotor activity. In addition, an open arm index was calculated for time spent (**F:** F_(2, 56)_=11.19, p<0.0001) and distance travelled (**H:** F_(2, 56)_= 10.99, p<0.0001) (full details in *Methods*). Values are the means ± SEM. One-way ANOVA. Group comparisons were made using Tukey’s multiple comparisons test and adjusted p-values are represented as: *p<0.05, **p<0.01, ***p<0.005.

**Figure 2.**
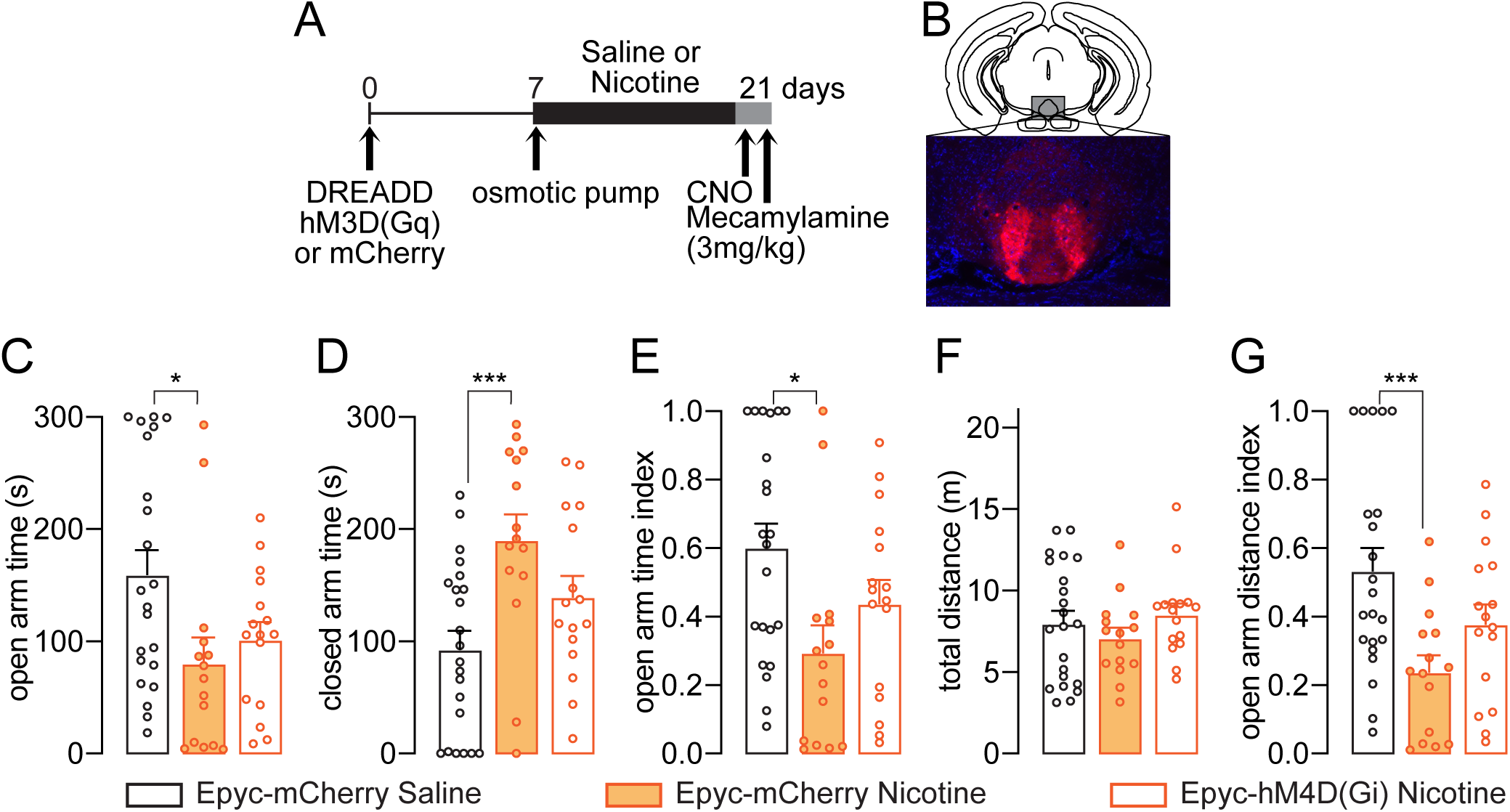
Inhibition of Epyc-IPN neurons fails to alter anxiety-like behaviors in nicotine exposed mice after precipitated withdrawal. **A)** Timeline of the experimental design. Experimental groups are as follows: mCherry Saline (N=22), mCherry Nicotine (N=15) and hM4D(Gi) Nicotine (N=16). **B)** Schematic diagram of virus injection target and representative image of virus expression in the IPN of Epyc-cre male and female adult mice. **C-G)** Elevated plus maze. Maze exploration was assessed by evaluating time spent in the open (**C:** F_(2, 50)_=4.210, p=0.0204) and closed (**D:** F_(2, 50)_=6.895, p=0.0023) arms as well as the total distance travelled in the maze (**F:** F_(2, 50)_=0.8902, p=0.4170) as a measure of locomotor activity. In addition, an open arm index was calculated for time spent (**E:** F_(2, 50)_=4.663, p=0.0139) and distance travelled (**G:** F_(2, 50)_=6.130, p=0.0042) (full details in *Methods*). Values are the means ± SEM. One-way ANOVA. Group comparisons were made using Tukey’s multiple comparisons test and adjusted p-values are represented as: *p<0.05, **p<0.01, ***p<0.005.

Alterations in physical behaviors of the sensorimotor domain such as grooming, scratching, twitching and other stereotyped actions have been reported in the literature as somatic manifestations of the withdrawal syndrome in rodents (Malin et al., 1992; Malin et al., 1994; Damaj et al., 2003). To evaluate this aspect of the syndrome, immediately after the EPM test (**Figures 1,2**) we placed the animals in an open field and recorded their behavior for 15 minutes. We observed two types of behaviors: 1) grooming-related and 2) shakes & twitches (detailed description in *Methods*) and found that neither the number of these categories of events (18.41±2.31 vs. 23.67±3.11, **Figure 3A**; 8.73±1.65 vs. 8.73±1.11, **Figure 3C**; 28.30±3.58 vs. 35.90±5.15, **Figure 3F**; 6.40±1.03 vs. 5.05±0.65, **Figure 3H**) nor the time spent engaged in these behaviors (46.18±5.55 s vs. 46.17±4.86 s, **Figure 3B**; 1.45±0.26 s vs. 1.519±0.22 s, **Figure 3D**; 44.80±4.81 s vs. 54.23±6.32 s, **Figure 3G**; 1.00±0.16 s vs. 0.78±0.88 s, **Figure 3I**) was affected by nicotine exposure and precipitated withdrawal compared to saline mCherry controls. Moreover, inhibition of Amigo1 or Epyc-IPN neurons failed to alter these behaviors (23.67±3.11 vs. 25.19±2.01, **Figure 3A**; 46.17±4.86 s vs. 31.81±4.29 s, **Figure 3B**; 8.73±1.11 vs. 8.44±0.90 **Figure 3C**; 1.52±0.22 s vs.1.47±0.18 s, **Figure 3D**; 35.90±5.15 vs. 31.21±4.26, **Figure 3F**; 54.23±6.32 s vs. 37.66±4.88 s, **Figure 3G**; 5.05±0.65 vs. 6.68±1.08, **Figure 3H**; 0.78±0.88 s vs. 1.04±0.17 s, **Figure 3I**). No change was observed either in the distance travelled in the maze by all the experimental groups (26.84±2.07 m, 25.02±1.66 m, 25.95±162 m, **Figure 3E**; 31.37±2.93 m, 31.41±3.32 m, 29.41±4.60 m, **Figure 3J**). These findings reveal that the present model of nicotine exposure and withdrawal does not elicit any alteration in physical signs and that reducing the activity of Epyc or Amigo1-IPN neurons is also insufficient to effect changes in the somatic domain.

**Figure 3.**
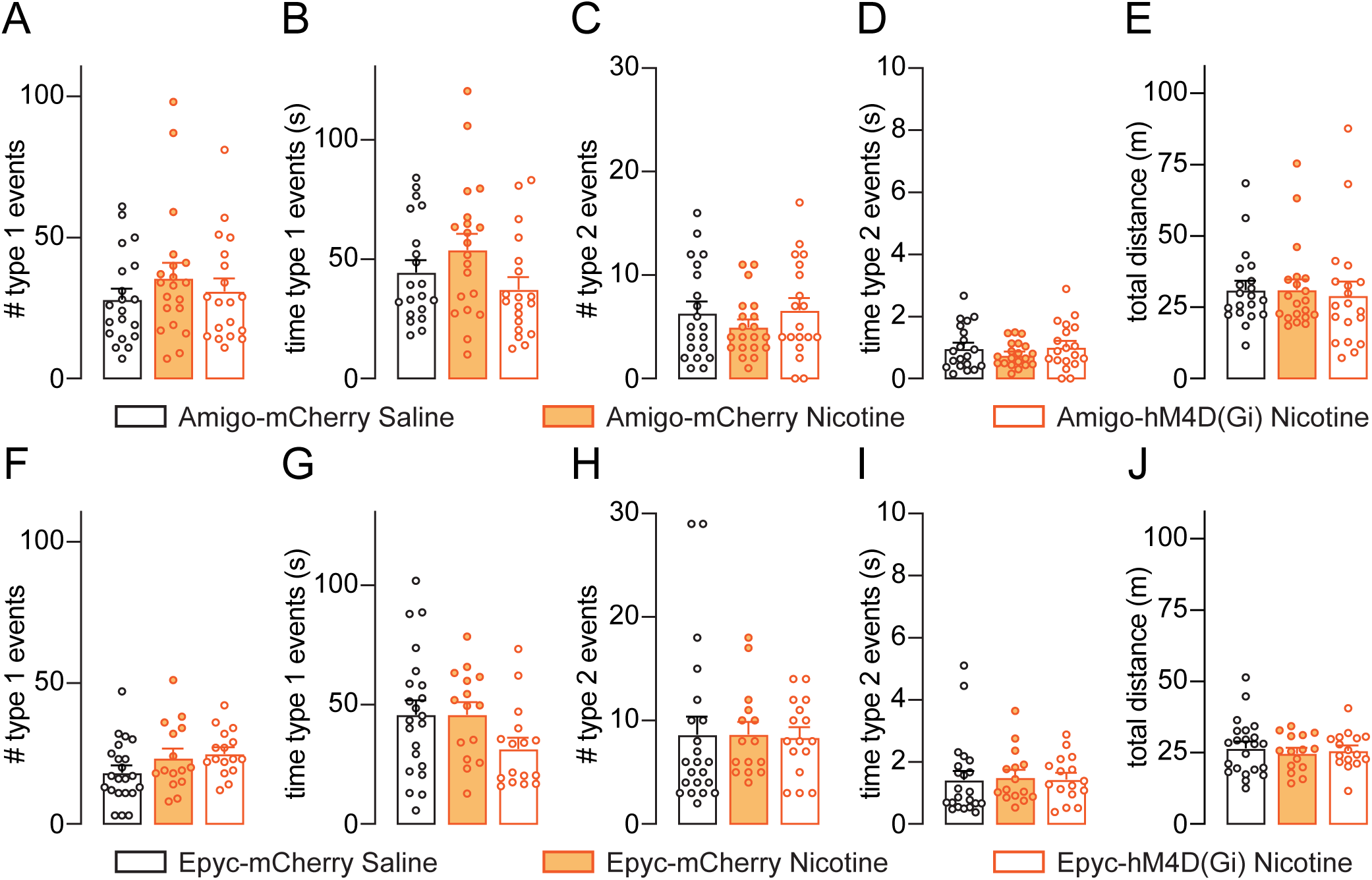
Somatic behaviors are unaffected by nicotine withdrawal and inhibition of Amigo1-or Epyc-IPN neurons. Somatic signs of withdrawal were grouped in two types: 1) grooming-related (**A-B**, **F-G**) and 2) shakes & twitches (**C-D**, **H-I**) (full description in *Methods*). The total distance travelled in the arena was included as a measure of locomotor activity (**E, J**). Values are represented as means ± SEM and analyzed using one-way ANOVA. No behavioral change was apparent between nicotine exposed and nicotine naïve Amigo1-mCherry animals with respect to the number (**A:** F_(2, 56)_=0.7782, p=0.4641; **C:** F_(2, 56)_=0.8737, p=0.4230) and time (**B:** F_(2, 56)_=2.356, p=0.1041**; D:** F_(2, 56)_=0.9727, p=0.3843) spent engaged in these behaviors or the distance travelled in the arena (**E:** F_(2, 56)_=0.09621, p=0.9084). Inhibition of Amigo1-IPN neurons in nicotine exposed animals failed to alter any of these behaviors with respect to their mCherry counterparts. Similarly, no difference in the number (**F:** F_(2, 50)_= 2.230, p=0.1181**; H:** F_(2, 50)_=0.01384, p=0.9863) or time (**G:** F_(2, 50)_=2.442, p=0.0973; **I:** F_(2, 50)_= 0.02361, p=0.9767) of somatic signs was observed between nicotine exposed and nicotine naïve Epyc-mCherry or Epyc-hM4D(Gi) Nicotine animals. These groups also presented similar distance travelled in the arena (**J**: F_(2, 50)_= 0.2320, p=0.7938).

To determine the extent to which Amigo1 cells regulate anxiety-like behaviors we next asked whether stimulation of Amigo1-IPN neurons is sufficient to elicit such behaviors in the absence of nicotine exposure (**Figure 4A** for timeline). For that purpose, Amigo1-cre animals were infected in their IPN by AAV9-hSyn-DIO-hM3D(Gq)-mCherry, which delivers an excitatory hM3D(Gq) cre-dependent DREADD fused with mCherry, or AAV-DIO-mCherry as a control. To test the effect of the DREADD on IPN neuron activity, we recorded hM3D(Gq)-mCherry IPN neurons using patch clamp slice electrophysiology as described previously. Here, we found that after focal delivery of CNO and equilibration, Amigo1 neurons showed marked increase in action potential frequency, necessitating a negative DC current injection to determine a voltage at rheobase, which shifted from -56.1 ± 2.6 mV to 64.1 ± 3.9 mV upon CNO exposure (p=0.013, **Figure 4C** *right*), indicating that CNO could effectively potentiate excitability in hM3D(Gq) expressing Amigo1-cre neurons. Conversely, we recorded no significant change to passive properties measured by resting potential, which was -57.8 ± 2.7 mV before and -55.6 ± 3.0 mV after CNO exposure (p=0.22, left **Figure 4C** *left*). We then quantified affective behaviors in Amigo1-hM3D(Gq) as described previously. Here, we found that stimulation of Amigo1-IPN neurons was sufficient to reduce the exploration of the open arms of the EPM in nicotine-naïve mice (**Figure 4D-F**). This is evidenced by the pattern of reduced time spent (time 178.40±23.89 s vs. 60.60±17.81 s, p=0.0004, **Figure 4D**; index 0.65±0.08 vs. 0.24±0.07, p=0.0006, **Figure 4F**) and distance travelled (index 0.60±0.08 vs. 0.24±0.07, p=0.0018, **Figure 4H**), in the open arms of the maze compared to saline controls which is indistinguishable from the open arm behavior of non-DREADD mice undergoing nicotine withdrawal (time 60.60±17.81 s vs 68.23±13.98 s, p=0.9609 **Figure 4D**; index 0.24±0.07 vs. 0.28±0.06, p=0.9343, **Figure 4F**; distance index 0.24±0.07 vs 0.26±0.06, p=0.9892, **Figure 4H**). This pattern is still clear when accounting for the increased locomotor activity of these animals (total distance travelled 7.21±0.72 m vs. 14.58±1.26 m, p<0.0001, **Figure 4G**) as shown by the open arm distance index (**Figure 4H**). Taken together these results show that manipulating activity of Amigo1-IPN neurons is sufficient to regulate anxiety-like behaviors in mice undergoing nicotine withdrawal as well as mice who are nicotine-naive.

**Figure 4.**
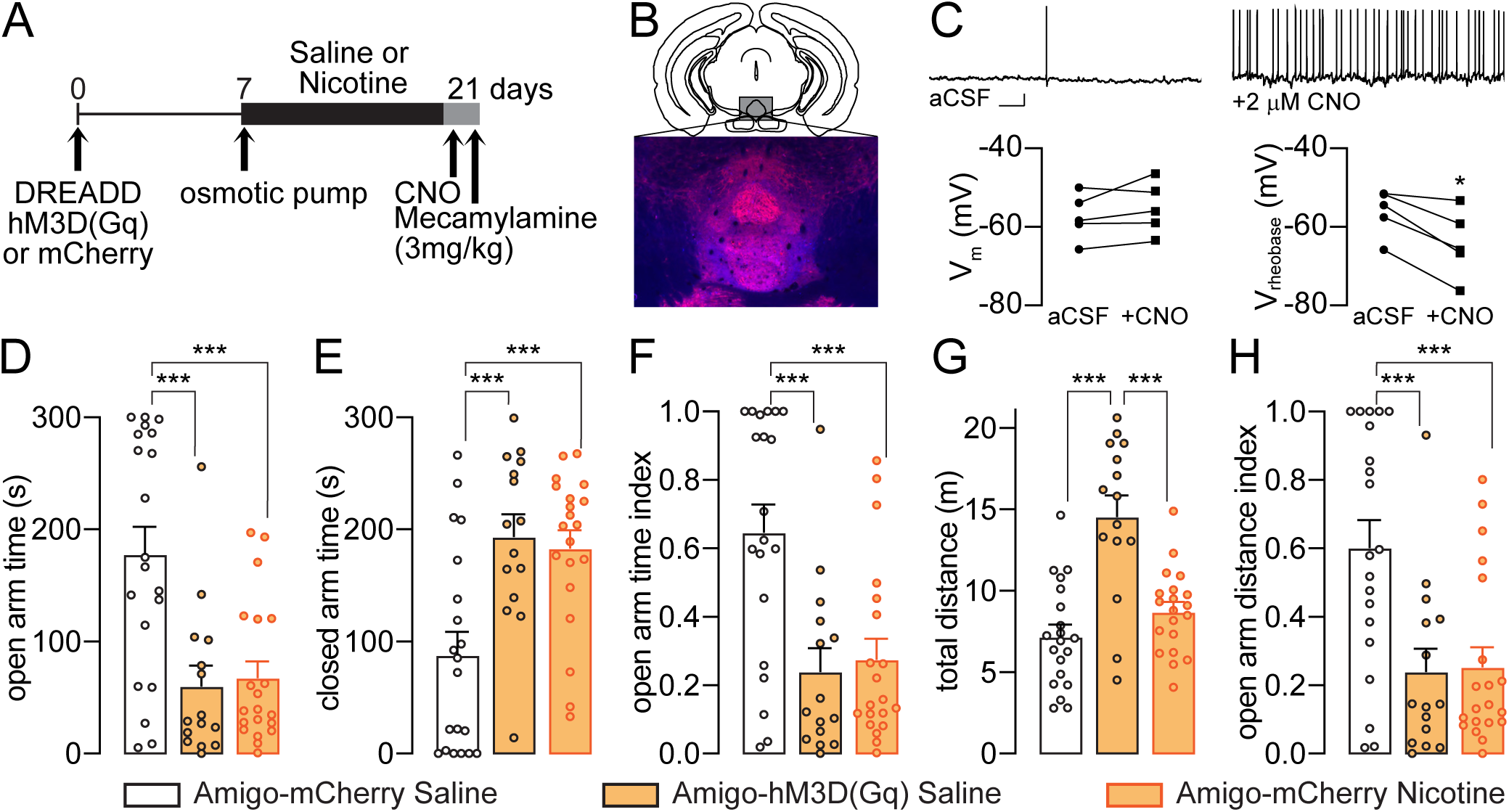
Stimulation of Amigo1-IPN neurons is sufficient to elicit anxiety-like behaviors in nicotine-naïve mice. **A)** Timeline of the experimental design. Experimental groups are as follows: mCherry Saline (N=20), mCherry Nicotine (N=20), hM3D(Gq) Saline (N=15). **B)** Schematic diagram of virus injection target and representative image of virus expression in the IPN of Amigo1-cre male and female adult mice. **C)***Top*: Sample traces of action potentials from Amigo1-cre cells expressing hM3D(Gq) in response to aCSF alone or supplemented with 2μM CNO. Scale bar indicates 10mV and 500ms and applies to all traces. *Left*: Resting membrane potential before and after CNO application (n=5, t-test). *Right*: A negative shift in the rheobase for action potential voltage (f<0.1 Hz) is observed following CNO application (n=5, t-test). **D-H)** Elevated plus maze. Maze exploration was assessed by evaluating time spent in the open (**D:** F_(2, 52)_=11.93, p<0.0001) and closed (**E:** F_(2, 52)_=10.18, p=0.0002) arms as well as the total distance travelled in the maze (**G:** F_(2, 52)_=19.94, p<0.0001) as a measure of locomotor activity. In addition, an open arm index was calculated for time spent (**F:** F_(2, 52)_=10.71, p=0.0001) and distance travelled (**H:** F_(2, 52)_=9.392, p=0.0003) (full details in *Methods*). Values are the means ± SEM. One-way ANOVA. Group comparisons were made using Tukey’s multiple comparisons test and adjusted p-values are represented as: *p<0.05, **p<0.01, ***p<0.005.

Next, we tested whether Amigo1-IPN stimulation would affect somatic signs of withdrawal, as described previously. Mice were placed in an open field and observed for somatic signs of nicotine withdrawal, as described (**Figure 5**). In contrast with our observations from inhibitory DREADD, stimulation of Amigo1-IPN neurons significantly reduced the time spent and the total number of both grooming-related behaviors (type 1, 28.30±3.58 s vs.11.13±2.60 s, p=0.0009, **Figure 5A**; 44.80±4.81 vs. 10.59±1.65 events, p<0.0001, **Figure 5B**) and shakes & twitches (type 2, 1.00±0.16 s vs. 0.21±0.05 s, p=0.0002, **Figure C**; 6.40±1.03 vs. 1.27±0.28 events, p=0.0002, **Figure 5D**) compared with nicotine-naïve controls suggesting that Amigo1-IPN neurons are part of the circuit responsible for these behaviors under basal conditions. This manipulation also resulted in an increase of the locomotor activity evidenced by an increase in the total distance travelled in the arena (31.37±2.93 m vs. 50.25±7.14 m, p=0.0112, **Figure 5E**). Taken together, these results indicate that Amigo1-IPN neurons are differentially involved in two behavioral phenotypes typically associated with withdrawal after nicotine exposure such that their stimulation elicits clear changes in opposite directions (i.e. increased anxiety-like behavior and reduced somatic signs).

**Figure 5.**
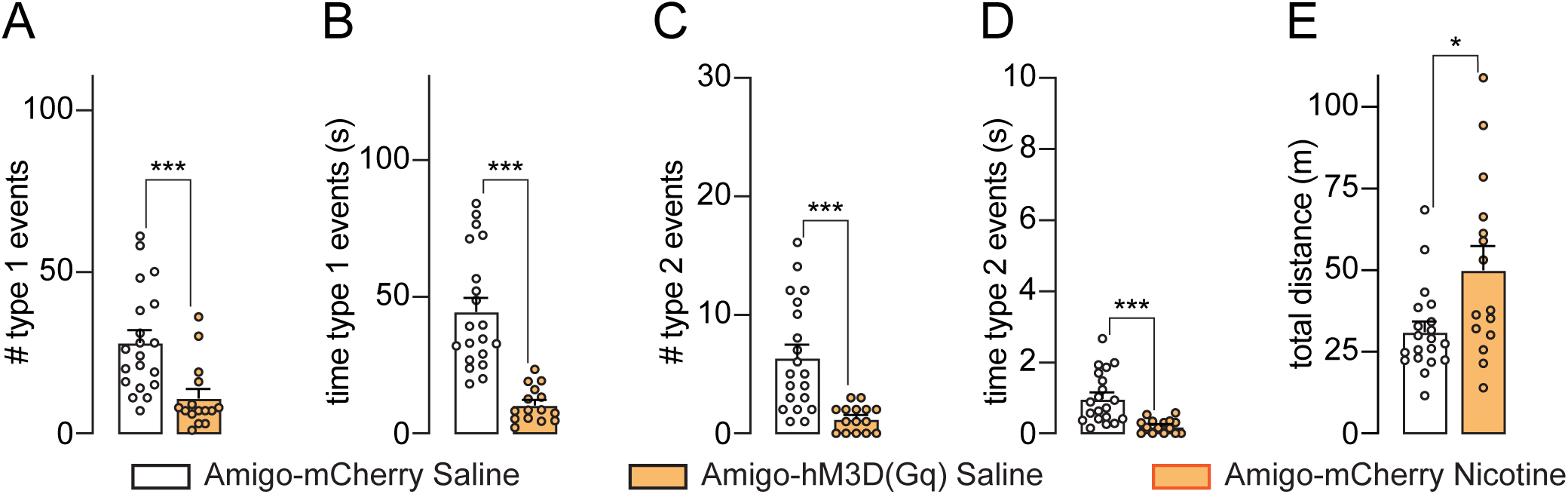
Stimulation of Amigo1-IPN neurons reduces somatic behaviors in nicotine-naïve mice while increasing overall locomotor activity. Somatic signs of withdrawal were grouped in two types: 1) grooming-related (**A, B**) and 2) shakes & twitches (**C, D**) (full description in *Methods*). The total distance travelled in the arena was assessed as a measure of locomotor activity (**E**). Values are the means ± SEM. Group comparisons were made using a two-tailed unpaired t-test (p-values are represented as: *p<0.05, **p<0.01, ***p<0.005). The number of events (**A:** p=0.0009**; C:** p=0.0002) and time (**B:** p<0.0001; **D:** p=0.0002) spent engaged in somatic behaviors was significantly reduced following stimulation of Amigo1-IPN neurons. Notably, the total distance travelled in the arena was in fact increased with respect to their mCherry counterparts (**J:** p=0.0112).

## Discussion

The IPN plays a crucial role in the somatic and affective behaviors that characterize nicotine withdrawal syndrome (Salas et al., 2009; Zhao-Shea et al., 2013; Klenowski et al., 2022). Several lines of evidence had indicated that IPN is heterogeneous and contains different cell types marked by different calcium binding proteins, neuropeptides, and nAChR types (Hsu et al., 2013; Zhao-Shea et al., 2013; Shih et al., 2014). However, Ables and colleagues provided the first description of non-overlapping neuronal subpopulations with separate behavioral outcomes (Ables et al., 2017). Here, we leveraged this distinction to independently target Amigo1 or Epyc neuronal subpopulations within the IPN. We found that Amigo1-IPN cells are essential in regulating anxiety-like behavior in mice navigating an EPM. By effectively modulating the activity of this subset of GABAergic IPN neurons we could prevent or induce the characteristic affective phenotype. Thus, activity within the subpopulation of Amigo1-IPN neurons is both necessary and sufficient for anxiety-like behavior in mice. This agrees with the evidence establishing the IPN as a mediator of negative emotional states, such as fear and anxiety both in basal and nicotine-withdrawal conditions (Molas et al., 2017). In contrast, we did not find any effect of chemogenetically manipulating a separate pool of α5 nAChR-expressing GABAergic neurons in the IPN, marked by expression of Epyc, indicating that the cellular heterogeneity in this nucleus carries forward to the role of each neuron type in regulating behavior. The main source of glutamatergic and cholinergic innervation to both GABAergic subpopulations in the IPN comes from the mHb (Ren et al., 2011; Frahm et al., 2015; Ables et al., 2017) and is reciprocally connected to brain structures associated with anxiety such as the laterodorsal tegmentum and the serotonergic Raphe nuclei. (Lima et al., 2017; Quina et al., 2017). The ventral tegmental area could be another key input to the IPN, but this projection remains controversial (Quina et al., 2017; DeGroot et al., 2020). Taken together, these findings implicate suggest that both the internal and external functional connectivity of the IPN require further characterization to fully elucidate how it drives the nicotine withdrawal phenotype.

In contrast to studies linking nicotine and affective behavior, past studies of somatic manifestations of nicotine withdrawal are more variable in the behaviors evaluated and how they are assessed. Although many experimenters use a combined withdrawal score, others report that nicotine withdrawal does not elicit an increase in all somatic signs analyzed (Malin et al., 1992; Zhao-Shea et al., 2013; Klenowski et al., 2022). Recognizing the challenges of assessing this aspect of the syndrome, we implemented a scoring system to precisely record the time and duration of somatic signs and compare results among independent scorers to confirm that our criteria to identify the relevant behaviors (i.e. grooming, scratching, genital licking, shaking, twitching or jumping) was consistent. However, after thorough and detailed analysis we observed that under our baseline conditions nicotine-exposed animals did not exhibit a statistically significant change in the number or timing of somatic signs following precipitation of withdrawal. This may reflect the specific conditions of our experiments and may also be partly due to not observing any significant prevalence of behaviors that reportedly changed in other studies (e.g. teeth chattering, gasps, backing). We omitted digging behaviors (by not providing any substrate in the arena) as this behavior likely overlaps with the affective phenotype, given the established association of marble-burying with anxiogenesis during nicotine withdrawal (Zhao-Shea et al., 2015; Chellian et al., 2021). Taken together, our results indicate that affective and somatic signs are separable components of the nicotine withdrawal phenotype that are dissociable at the behavioral level.

Chemogenetic stimulation of Amigo1-IPN neuron activity reduced the number and time of somatic signs in nicotine-naïve mice, an effect that is opposite to the nicotine withdrawal phenotype. Manipulation of Amigo1-IPN thus does not mimic the overall withdrawal phenotype. Instead, these data suggest that Amigo1-IPN neurons may lie at the intersection of two related but different circuits underlying the physical and affective components of the withdrawal phenotype which may integrate different relevant afferent information (e.g. from the mHb, medial raphe nucleus or putative dopaminergic inputs) (Lima et al., 2017; Quina et al., 2017; DeGroot et al., 2020). Other manipulation to IPN neuron activity such as optogenetic stimulation of GAD2+ IPN differentially impacts these two behavioral domains by increasing somatic signs while not eliciting any changes in anxiety-like behavior (Zhao-Shea et al., 2013). However, a similar manipulation has been shown to reduce anxiety-like behavior in animals undergoing spontaneous withdrawal (Klenowski et al., 2022). In addition to differences associated with different models of nicotine exposure (constant subcutaneous delivery or intermittent oral consumption) and withdrawal (precipitated or spontaneous), our findings reveal the need for subpopulation-specific studies of the mechanisms underlying nicotine withdrawal.

By separately manipulating Amigo1 and Epyc GABAergic populations within the IPN, we have demonstrated that somatic and affective signs of withdrawal are separable components of the phenotype at the behavioral level. Future studies of how nicotine exposure drives plasticity in the different IPN synapses will further differentiate these phenotypes. In this sense, an important future objective is to identify the different circuitry engaged up and downstream the Amigo1 subpopulation of IPN neurons, both in the context of understanding and treating nicotine withdrawal and in leveraging the contribution of this circuit to general anxiety phenotypes.

## Acknowledgements

Supported by Institutional funds from the University of Illinois at Chicago and NIH grant R00-DA041500 to CJP.

